# Comparative Phylogenomics and Phylotranscriptomics Provide Insights into the Genetic Complexity of Nitrogen Fixing Root Nodule Symbiosis

**DOI:** 10.1101/2023.04.03.535273

**Authors:** Yu Zhang, Yuan Fu, Wenfei Xian, Xiuli Li, Yong Feng, Fengjiao Bu, Yan Shi, Shiyu Chen, Robin van Velzen, Alison M. Berry, Marco G. Salgado, Hui Liu, Tingshuang Yi, Pascale Fournier, Nicole Alloisio, Petar Pujic, Hasna Boubakri, M. Eric Schranz, Pierre-Marc Delaux, Gane Ka-shu Wong, Valerie Hocher, Sergio Svistoonoff, Hassen Gherbi, Ertao Wang, Wouter Kohlen, Luis G. Wall, Martin Parniske, Katharina Pawlowski, Normand Philippe, Jeffrey J. Doyle, Shifeng Cheng

## Abstract

Plant root nodule symbiosis (RNS) with mutualistic nitrogen-fixing bacteria is restricted to a single clade of angiosperms, the Nitrogen-Fixing Nodulation Clade (NFNC), and is best understood in the legume family. It is widely accepted that nodulation originated through the assembly of modules recruited from existing functions, such as mycorrhizal symbiosis, polar growth, and lateral root development. Because nodulating species are scattered within the NFNC, the number of times nodulation has evolved or has been lost has been a matter of considerable speculation. This interesting evolutionary question has practical implications concerning the ease with which nodulation might be engineered in non-nodulating crop plants. Nodulating species share many commonalities, due either to divergence from a common ancestor over 100 million years ago or to convergence or deep homology following independent origins over that same time period. In either case, comparative analyses of diverse nodulation syndromes can provide insights into constraints on nodulation—what must be acquired or cannot be lost for a functional symbiosis—and what the latitude is for variation in the symbiosis. However, much remains to be learned about nodulation, especially outside of legumes. Here we present new information across the spectrum of nodulating groups. We find no evidence for convergence at the level of amino acid residues or gene family expansion across the NFNC. Our phylogenomic analyses further emphasize the uniqueness of the transcription factor, NIN, as a master regulator of nodulation, and identify key mutations affecting its function across the NFNC. We find that nodulation genes are over-represented among orthologous gene groups (OGs) present in the NFNC common ancestor, but that lineage-specific OGs play major roles in nodulation. We identified over 900,000 conserved noncoding elements (CNEs), of which over 300,000 were unique to NFNC species. A significant proportion of these are associated with nodulation-related genes and thus are candidates for transcriptional regulators.

## Introduction

Nitrogen is one of the main nutrient elements indispensable for plant growth and development, but it is not accessible directly to plants without the help of the nitrogenase-containing nitrogen-fixing bacteria (Geurts et al., 2016; Mathesius, 2022). Some plants can obtain ammonium quite effectively by accommodating nitrogen-fixing bacteria in a specialized root organ, the symbiotic nodule, allowing them to grow even in nitrogen-poor soils. How nodulation evolved is a fascinating question in its own right, but the ability of nodulating plants to acquire nitrogen without exogenous fertilizer makes understanding how plants recruited and assembled the diverse components required for functioning nodules a topic of agronomic, economic and ecological importance.

The best-known nodulating species belong to the legume family (Fabaceae; e.g., soybean, pea, alfalfa) in the flowering plant order Fabales, but nodulation also occurs in three other orders (Fagales, Rosales, Cucurbitales). There is rich genetic, phenotypic, and eco-adaptive diversity among nodulating plants across the four orders, including biogeographic distribution, nodule ontogeny, infection mode and formation of intracellular endosymbiosis across the different lineages (Shen et al., 2020). Most notable is the diversity of the microsymbionts: Legumes (Fabales) and *Parasponia* (Cannabaceae, Rosales) associate with a group of Gram-negative nitrogen fixing soil bacteria collectively called rhizobia (Ardley and Sprent, 2021; Sprent et al., 2017), whereas the remaining nodulating species from Fagales, Rosales, and Cucurbitales engage with actinobacteria of the genus *Frankia,* and are termed actinorhizal plants. This diversity, coupled with the fact that the four orders were distantly related in pre-phylogenetic classification systems, suggested that there could be many paths to nodulation.

A major result of early molecular phylogenetic studies was the placement of these four orders in a monophyletic “Nitrogen Fixing Nodulation Clade” (NFNC) within the large Rosid clade of angiosperms (Soltis et al., 1995). This led Soltis et al. (1995) to hypothesize that a “predisposition” for nodulation evolved in the most recent common ancestor (MRCA) of the NFNC over 100 million years ago (MYA) that was either the nodulation synnovation (Donoghue and Sanderson, 2015) itself (single origin hypothesis; **Scenario I**) or an unknown precursor trait that conferred a propensity for a nodulation syndrome (Sinnott Armstrong et al., 2022) to evolve independently several different times convergently, in some cases many millions of years after the NFNC MRCA (multiple origins model; **Scenario II**) (Soltis et al., 1995). If a constraining predisposition trait could be identified, Scenario II would provide more hope for engineering the full nodulation syndrome in non-NFNC species.

Nodulating species are found in only 10 out of the 28 plant families within the NFNC, and are rare in most of these families; even in the Leguminosae, although in some large clades nearly all species nodulate, much of the phylogenetic diversity of the legume family is non-nodulating (Doyle, 2011). Therefore, Scenario I requires massive parallel losses of nodulation across the NFNC, and because of this both intuitive reasoning and formal modeling studies have long favored the precursor/multiple origins Scenario II model (Battenberg et al., 2018; Doyle, 2011; Kates et al., 2022; Li et al., 2015; Werner et al., 2014), Recently, however, acceptance of Scenario I has been driven by two studies reporting genomic evidence consistent with non-nodulating NFNC species having lost the ability to nodulate rather than lacking it fundamentally (Griesmann et al., 2018; Van Velzen et al., 2018). Moreover, the current cessation of nodulation in many unrelated species can be explained by global reduction in the benefit of nitrogen relative to the cost of carbon as atmospheric CO_2_ levels decreased over the last ∼100 MY (van Velzen et al., 2019). Acceptance of Scenario I by many in the nodulation community is also fueled by the fact that after nearly three decades of intense searching, the precursor trait required by Scenario II has yet to be identified, though some candidates have been suggested (e.g., Mergaert et al., 2020; Miri et al., 2016; Soyano and Hayashi, 2014).

Root nodule symbiosis is not a single trait, but a complex association involving a number of integrated but independent genetic processes, including intracellular recognition and signaling (Fournier et al., 2018), nodule organogenesis, nitrogen responses, trophic exchanges and bacteria accommodation (Mathesius, 2022; Mergaert *et al*., 2020; Soyano and Hayashi, 2014; Soyano et al., 2021). In both scenarios, key components of the syndrome were recruited from other phenomena (e.g., Doyle, 1994; 2016), notably mycorrhizal signaling (Markmann and Parniske, 2009; Wang et al., 2022), but also from other pre-existing processes, such as elements of the lateral root development program (Soyano et al., 2021). In Scenario I the diversity of nodulation is due to divergence of homologous traits over more than 100 MY, whereas in Scenario II similarities in non-homologous nodules and the processes by which they are formed are due to convergence, including convergent recruitment of the same traits leading to “deep homology” (Shubin et al., 2009). Regardless, however, commonalities shared by unrelated nodulating species represent potential constraints on the process of nodulation—elements that are required for nodulation and are present in all nodulation symbioses either because they are features inherited from the only originator of nodulation, or because they had to be recruited convergently to build an effective nodulation symbiosis.

With a clearer understanding of the comparative biology of nodulation across the NFNC, it may be possible to identify the features that make the NFNC unique among plants in generating numerous lineages containing species that fix nitrogen in symbiotic partnership with bacteria that otherwise would be recognized as enemies (Parniske, 2018). Greater knowledge of non-nodulating NFNC taxa would also be useful (e.g., Tokumoto et al., 2020), particularly lineages in which nodulation has clearly been lost (Billault-Penneteau et al., 2019).

In this study, we have filled in some gaps with new genomes and transcriptomes and, together with the public dataset, we performed phylogenomic and phylotranscriptomic analyses within the NFNC. We explored genomic variation, gene expression changes and regulatory sequences that are candidates for driving the origin and diversification of plant nodulation.

## Results

### New genome and transcriptome sequences from NFNC species

Actinorhizal plants have been underrepresented relative to Fabaceae in availability of genomic and transcriptomic data, which is unfortunate given their phylogenetic diversity. To fill in this gap, we added three plant species to our previous sampling (Griesmann et al., 2018): 1) a non-nodulating member of Fagales (*Fagus sylvatica*; also recently sequenced by Mishra et al., 2018, 2021 (Mishra et al., 2018; Mishra et al., 2021)) to complement *Juglans regia*; 2) the non-nodulating *Dryas octopetala* to complement nodulating *Dryas drummondii* (Billault-Penneteau *et al*., 2019) to form a contrasting comparison pair within the same Rosales genus; and 3) the nodulating plant *Purshia tridentata* (also from the Rosales (**Figure 1, Table S2**)).

**Figure 1.**
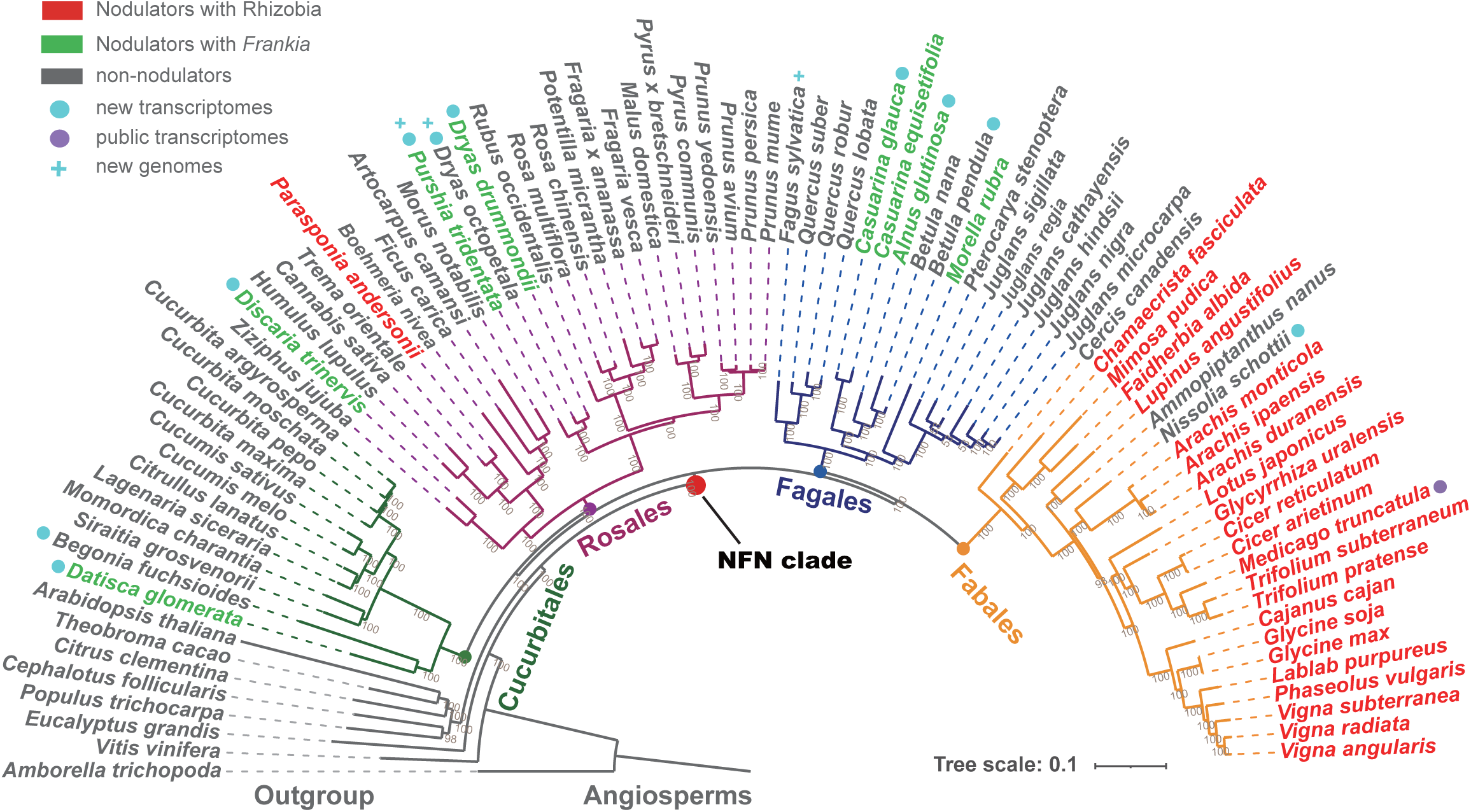
Genome and transcriptome dataset used in this study was illustratively summarized in a phylogenetic framework. The 88 genomes and representative transcriptomes used in this study are shown in a phylogenetic tree. The phylogenetic tree presented here for the 88 species, including three newly sequenced genomes, was using the reconstructed by the concatenated super matrix (47,473 amino acids) of 824 one to one orthologs that were present in all species. The tree was built by IQtree ML algorithm, with (Nguyen et al., 2015) JTT+F+R6 model (10,000 of bootstrap).

We sequenced these genomes using the traditional shotgun short-read sequencing technologies with hierarchical DNA-libraries of varied insert size (see Method), resulting in 361 Gb, 48 Gb, 130 Gb sequencing data for *Dryas octopetala* (257Mb – estimated genome size), *Fagus sylvatica* (497Mb), *Purshia tridentata* (244Mb). The two tree species are highly heterozygous, leading to a relatively fragmented genome assemblies even though more than 95% of the assemblies are covered by BUSCO core genes (**Table S2**). We obtained 28,191 (*Dryas octopetala*), 23,155 (*Purshia tridentata*), and 35,140 (*Fagus sylvatica*) gene models annotated from the assembled genomes for each species, respectively. By target gene sequence alignment and comparison, all of the 227 canonical legume symbiosis-related genes (Roy et al, 2020; van Velzen et al., 2018)(**Table S3**) were present in all three genomes. However, in the non-nodulating *D. octopetala*, the key nodulation regulator, NIN, is a pseudogene, having experienced a 69 nucleotide (21 amino acid) deletion relative to the intact gene in the congeneric nodulator, *D. drummondii* (**Figure S1**). This is in contrast to the other non-nodulator, *F. sylvatica*, in which NIN is intact as in other non-nodulating Fagales.

We also sequenced new transcriptomes from 19 phylodiverse nodulating and non-nodulating NFNC species (**Table S14**, and **Figure S7**), emphasizing root and nodule samples. We failed with some species (non-model) particularly in the nodule sample collection because of the difficulty in tissue culturing followed by RNA extraction (**Table S14, and Figure S7**). However, we generated high-quality RNA-seq dataset from at least one representative nodulating species from each of the four orders in the NFNC, including *Casuarina glauca* (Fagales), *Datisca glomerata* (Cucurbitales), *Purshia tridentata* (Rosales), and *M. truncatula* (Fabales) (**Figure 4A**), and built a complete treatment collection (B+/N-, B-/N+, B+/N+, B-/N-) in the mature root tissues for two closely-related comparison pairs: *Alnus* and *Betula* from Fagales (**Figure 4C**), *Datisca* and *Begonia* from Cucurbitales **(Figure S8)**. The pairwise correlations indicate a high-quality and consistent RNA-seq dataset within and between species (**Figure S7B**).

### Little evidence for convergence in proteins or gene family amplification

Proteins recruited for nodulation could evolve new, nodulation-specific amino acids. In Scenario II such changes could occur convergently, such that nodulating species would share sites not found in non-nodulating relatives. We implemented a test that Parker et al. (Parker et al., 2013) used to show convergence at a small number of amino acids in hundreds of genes across the genomes of echolocating mammals (bats and dolphins). We included genomes of 31 nodulating species and 49 closely related non-nodulating species within the NFNC, as well as 8 outgroup species (**Figure 1**, **Table S1**). The results show that the overwhelming majority of the tested orthologous genes (4,412/4,413), including all 54 symbiosis-related genes, generated gene trees consistently following the phylogenetic pattern of the species tree, as expected for orthologous genes (**Figure 2**, **Table S4**). We next examined the multiple sequence alignments for each of the 2,905 orthologous groups present only in all nodulating species (see Methods). We did not find any convergent sites shared only by all nodulating species. The Parker et al. (Parker *et al*., 2013) method is known to overpredict convergence (Thomas and Hahn, 2015), suggesting that our conclusion of a lack of convergence is robust.

**Figure 2.**
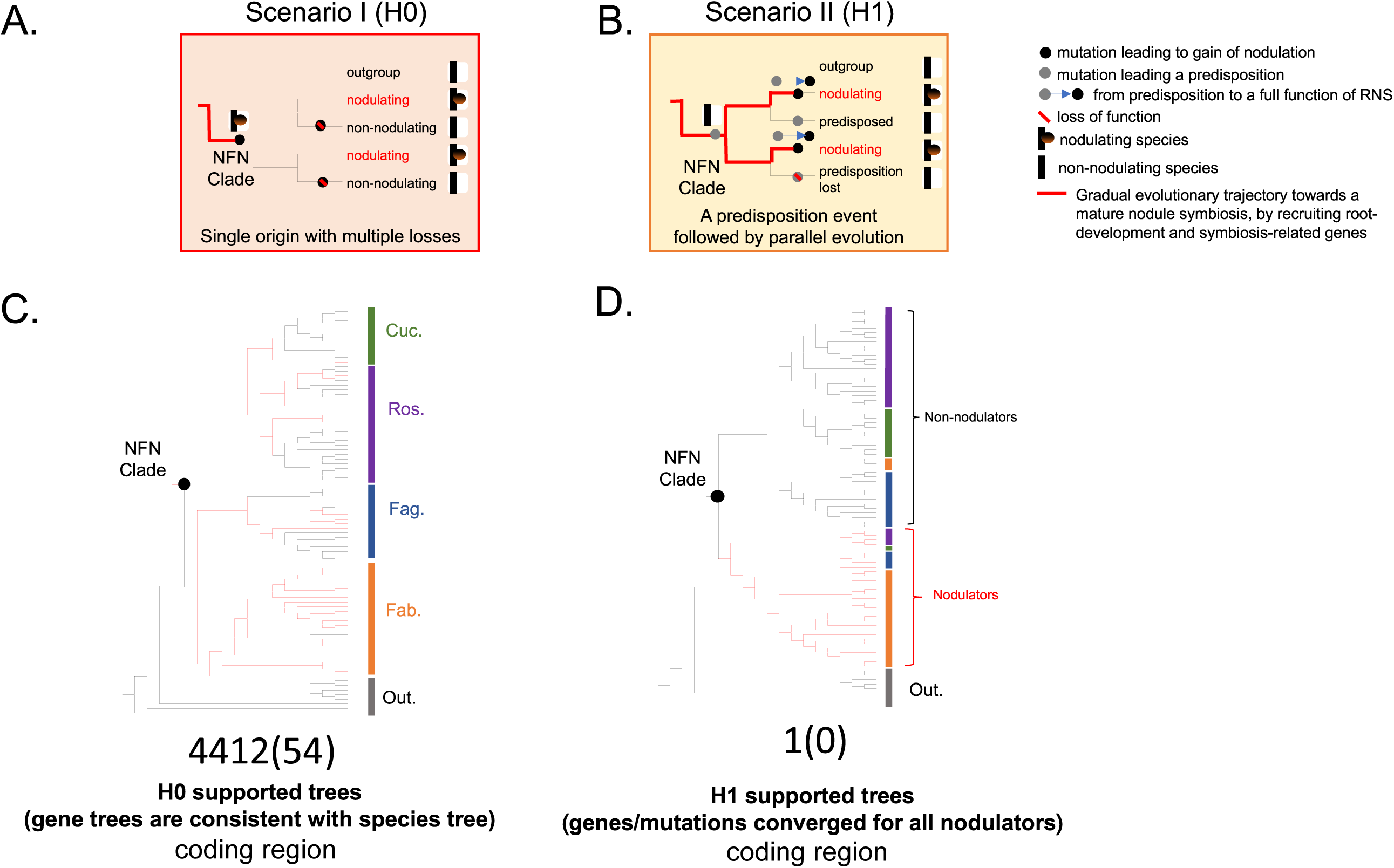
Phylogenetic inference of the two evolutionary scenarios on the origin of RNS based on thousands of gene orthogoups. Two evolutionary scenarios were established. **A.** Scenario I. single origin with multiple independent losses. This model predicts that novel gene(s) or mutations, which were sufficient to enable the establishment of a functional root nodule symbiosis, should have occurred at (exclusively in the ancestral status) or before (via a stepwise evolutionary journey long before) the common ancestor of the NFN clade. **B.** Scenario II. predisposition followed by parallel evolution, a common predisposition mutation (genomic innovation) took place in the common ancestor of the NFN clade, followed by a (series of) secondary decisive novel mutations that permitted a functional nodule symbiosis parallelly in different lineages. **C & D.** a tree topology inference method and an alignment method were implemented to identify what gene families follow either Scenario I (left) or Scenario II (right) proposed here, and to detect convergent signals (no convergent signal was treated as H0, as indicated in Scenario I) within orthologous sequences of nodulation species from different lineages. 4412 genes support H0 and 54 of them belong to symbiosis genes, but only one gene supports H1 and no symbiosis gene.

The origin of a complex novel phenomenon such as nodulation could involve the expansion of gene families; in the case of independent origins, expansion could involve some of the same families (Merényi et al., 2020). This was tested previously on a smaller subset of species (Griesmann et al., 2018; Zhao et al., 2021), revealing evidence of expansion that did not provide definitive support for either single or independent gain scenarios. We thus performed an association test on the significance of gene copy numbers between nodulating and non-nodulating species on a larger-scale (see Methods) (**Table S5-S7**). For the candidate orthogroups (53,657 OGs) detected across 88 species with gene annotations, we found that in total 96 gene families had experienced expansion in nodulating species compared with non-nodulating species (t-test p<0.01, difference of average of copy number > 1, **Tables S6, S7**), almost all of them being order-specific gene expansions that showed no convergence among different nodulating lineages. Furthermore, of these 96 expanding gene families, 8 encoded proteins were involved in the symbiosis (**Figure 3A, Table S7**) from legumes, which can be explained by the previous observations that complex ancestral polyploidization of the legumes might have led to the duplication of symbiosis genes (Koenen et al., 2021; Li et al., 2013; Zhao *et al*., 2021). Amplification of different gene families in diverse nodulating lineages could be due to refinement of nodulation after a single origin (Scenario I) or to independent origins of nodulation (Scenario II); more complex scenarios involving loss of family members could also be envisioned, particularly under Scenario I.

**Figure 3.**
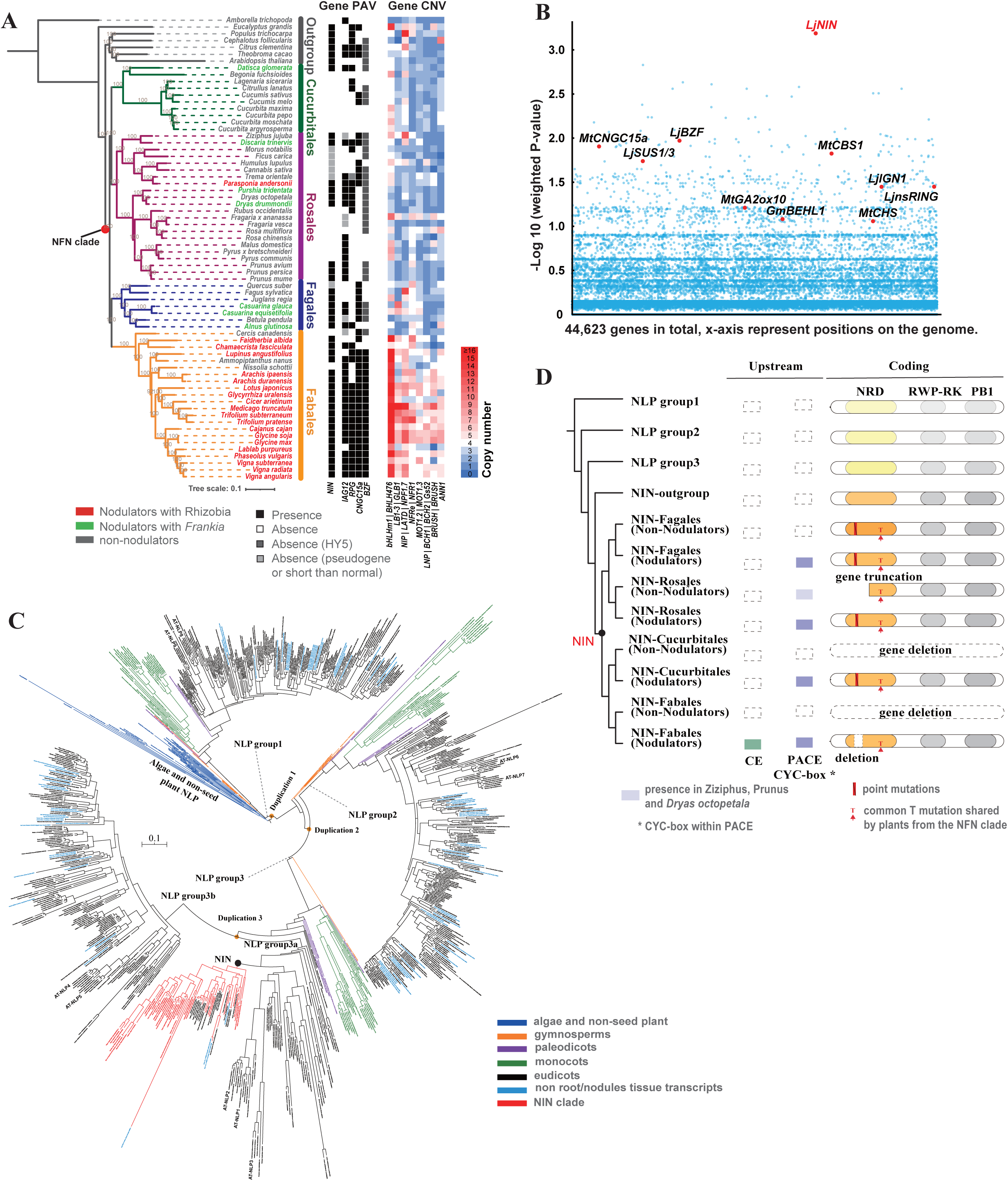
Gene presence and absence of symbiosis-related genes in the NFN clade, and the phylogenetic and structural analysis of NIN and NIN-like gene family. **A.** Occasional loss of symbiotic genes in a one-to-one ortholog detected by phylogenetic analysis in non-nodulating species and cases of gene families with putative contraction and expansion generated by orthofinder. **B.** Significance test of the association between gene presence and absence of the identified orthologous groups with the nodulation status between nodulating, non-nodulating and outgroup species. Fisher’s exact test was used to infer the association between presence of a target gene and the symbiosis status. Nodulation genes with weighted P-value > 1 were marked. **C.** The ML tree of the NIN-like protein family. The phylogenetic tree was built based on the multiple sequence alignment of NIN/NLP proteins using iqtree (JTT+I+G4, 1000 bootstraps) **D.** Schematic illustration and comparison of the protein and upstream sequence structural conservation, and variations between the different subgroups of NINs and NLPs, in which the evolutionary changes in the NRD region was highlighted.

To identify orthologs, to some extent, experiencing either loess or pseudogenization in non-nodulating species, we used a set of cutoffs to define ortholog presence in nodulating species (50% to 100% present, 6 conditions) and ortholog absence in non-nodulating species (50% to 100% absent, 6 conditions), resulting in 36 conditions (**Table S12**), this careful exploration is basically to avoid misinterpretation of the PAV results caused by varied genome data quality (in sequencing, assembly, annotation or alignment) derived from different studies. We finally defined the multiple losses of orthologous groups in non-nodulating species as that present in at least 70% of nodulating species and absent in at least 60% of non-nodulating species, resulting in the identification of 461 orthologs, including previously reported genes (*NIN* and *RPG*) and three additional symbiosis genes (*IAG12*, *CNGC15a*, *BZF*) through a gene-trait PAV association study (**Figure 3A, Table S11**). These identified genes play important roles in early signaling (*CNGC15a*), rhizobia/Frankia infection (*RPG*, *IAG12*), and nodule organogenesis (*NIN*, *BZF*).

### Sub-/neofunctionalization of the key nodulation-related gene *NIN* across the NFNC

We conducted genome-wide searches for the underlying genomic novelties from the protein-coding orthogroups that were specific to the NFNC, and then evaluated their association with nodulation. We integrated phylogenetic analysis for each gene family and Reciprocal Best Hit (RBH) search for each family member (**Table S8-12**) to detect fast-evolving genes, gene sub-clusters, or sequence neo-functionalization potentially specific to the common ancestor of the NFNC. This led to the identification of 37 orthologous gene families or sub-families that are absent in all outgroup taxa but present in at least 60% of NFNC species (**Table S8-10**). Among the 37 orthologous gene clusters, *NIN* stands out as the only one with functional validation of its involvement in nodule symbiosis (**Table S10**).

The 37 candidate genes were further evaluated individually for significant correlation between the number of species having the gene and whether or not that species nodulates. Again, *NIN* shows the most significant association with nodulating phenotype (**Figure 3B**, **Table S13**). We reconstructed the phylogenetic tree for all detected family members of NINs and NIN-like proteins (NLPs) in the genomes and transcriptome datasets (including 1KP transcriptomes) (**Figure 3C**), and obtained a tree that is largely consistent with that of a recent legume-focused study (Zhao *et al*., 2021) and other recent studies (e.g., Wu et al., 2022). The origin of the conserved *NLP-*like gene family that serves as a nitrate-responsive master regulator can be traced back to the common ancestor of green plants, subsequently experiencing at least three duplication events that resulted in four *NLP* subgroups. Consistent with previous observations (Liu and Bisseling, 2020), a duplication event within the *NLP* group 3 (*NLP3*) occurred early in the divergence of eudicots (Figure 3C, Duplication 3) and produced two paralogous clades, one of which includes *NIN* (in NFNC members) and its orthologs (in other eudicots).

To identify what sequence changes might have led to the functional transition from the as yet unknown ancestral function of *NLP* to the key nodulation roles of *NIN*, whether a single time or convergently, we identified nonsynonymous mutations in *NIN* that might have led to neo-functionalization specific to the NFNC. Two changes in *NIN* were identified that occurred in the common ancestor of the NFNC. The first is the loss or likely inactivation of the nitrate sensing motif in the nitrate responsive domain (NRD) of *NIN* conserved in non-NFNC species: all actinorhizal nodulating species examined here as well as *Parasponia andersonii* carry independent point mutations or small deletions in this motif in NIN compared with the NLP3 orthologs in the outgroup, and a larger deletion of this motif occurs in legume NINs (Suzuki et al., 2013) (**Figure 3D**, **Figure S3**). In *Arabidopsis*, phosphorylation of a serine residue (S205) within this nitrate sensing motif is indispensable for the relocation of the protein encoded by AtNLP7 from the cytoplasm to the nucleus, and the subsequent activation of downstream nitrate responsive genes (Liu and Bisseling, 2020). In contrast, in *Medicago*, NIN has lost the ability to sense nitrate and locates directly to the nucleus. The other NFNC-specific mutation occurred in the consensus position 363 in the NRD domain of NIN: an amino acid transition to threonine (363T event) (**Figure S3**). Although this 363T site is the only NIN-specific mutation that we found exclusively occurring in the most recent common ancestor of NFNC, it might not alone determine the ability to nodulate, consistent with our functional validation experiments (**Figure S4**). Much more remains to be learned about the structure and function of NIN in both nodulating and non-nodulating species.

### Phylotranscriptomic analysis reveals high conservation and an enhanced RNS gene expression network

Hundreds of protein-coding genes play roles during nodulation in *Medicago* (Tables S3; Roy et al., 2020), *Parasponia* (Van Velzen *et al*., 2018), and diverse actinorhizal nodulators (Battenberg et al., 2018; Diédhiou et al., 2014). However, much remains to be learned about the responses to nitrate or N_2_-fixing bacteria across different NFNC lineages. We thus performed a comparative phylotranscriptomic analysis for 227 functionally verified nodule symbiosis-related target genes (**Figure S7** and **Table S14**), which revealed that at least 6 symbiosis-related genes (*CHIT5, NF-YA1, CP6, NFH1, NIN, RSD*) covering almost all the stages of nodulation (and thus almost certainly recruited as separate modules once [Scenario I] or multiple times [Scenario II] in the evolution of nodulation), were commonly up-regulated in all the selected nodulating plant genomes (**Figure 5D**). 26 nodulation symbiosis-related orthologous genes were previously described as nodule upregulated genes in both *Parasponia* and *Medicago* (Van Velzen *et al*., 2018); following the same strategy, we detected 18, 19, and 14 *Medicago* nodule symbiosis-related genes upregulated in *Casuarina*, *Purshia* and *Datisca*, respectively. This pattern is consistent either with a retained function from a common nodulating ancestor (Scenario I) or convergent recruitment during independent origins of nodulation (Scenario II).

*NIN* orthologs were highly expressed almost exclusively in nodules of different nodulating species when inoculated with N_2_-fixing bacteria, but only under nitrogen-depleted conditions (**Figure 4B**), while NLP3 orthologs were relatively highly expressed in nitrogen depleted roots. Several genes were co-expressed with *NIN*: e.g., *GLB1* and *SST1* in nodules (**Figure 4A**), *bHLHm1* and *DWARF27* in roots (**Figure 4D)**. Phylogenetic analysis shows that the nodule-enhanced OGs from five gene families (*NIN, NF-YA1, NADH, NOOT, MCA8*) (Bu et al., 2020; Clavijo et al., 2015; Shen *et al*., 2020) were restricted in one-to-one orthologous gene clades (**Figure S6**), while the other genes had more complex profiles of expression and presented lineage specific upregulation of different paralogs.

**Figure 4.**
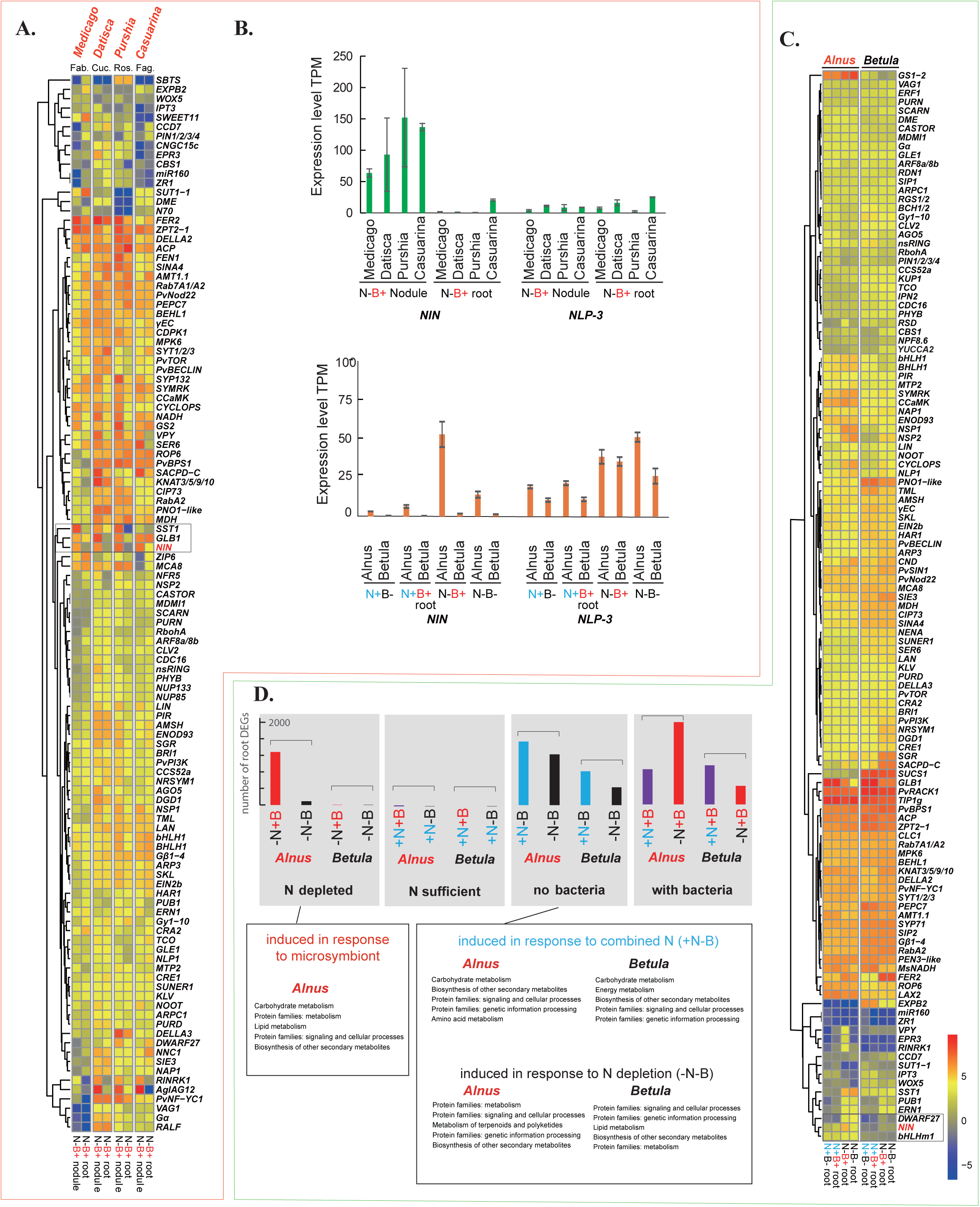
Differentially expressed genes responding to nitrate treatment and/or inoculation with N2-fixing bacteria within and between nodulating and non-nodulating plants in the NFN clade. **A.** Comparison of gene expression changes between root and nodule responding to the inoculation with N2-fixing bacteria (“N-B+” treatment) between the representative nodulating species (in red) within NFN clade (left). The hierarchical clustering of gene expression profile in roots/nodules was performed by R package hclust. The gene expression values (TPM) were calculated and compared based on the one to one orthologous symbiosis genes across species. (N, Nitrogen; B, N2-fixing Bacteria; +, with; -, without). **B.** Comparison of gene expression levels (TPM) of NIN and NLP-3 between nodules and roots for the selected plant species under different growth conditions. **C.** The detailed comparison between the ‘comparison pair’ of the nodulating plant *Alnus* and its closely-related species *Betula* in Fagales, under a various of treatments (N+B-, N+B+, N-B+, and N-). The gene expression values (TPM) were calculated and compared based on the one to one orthologous symbiosis genes across species. (N, Nitrogen; B, N2-fixing Bacteria; +, with; -, without). **D.** Summary of the differentially expressed genes (DEGs) in roots under different treatments with nitrate and microsymbiont. The bars represent the number of upregulated genes under a given growth condition compared with the other conditions. Blue, supplied with 5 mM KNO3; red, inoculated with Frankia; purple, supplied with 5 mM KNO3 and inoculated with *Frankia*; grey, supplied with neither KNO3 nor Frankia.

We then performed a genome-wide systematic evaluation of the connection between the conservation of OGs with the conservation in gene expression level across lineages. From the same four representative nodulating species described above we classified genes as follows: shared by all 4 orders; shared by species from 3 orders; shared by species from 2 orders where one species was from either Fabales or Fagales, and the other was from either Cucurbitales or Rosales; shared by species from two sister orders; or unique to a single species (**Figure 5A**). Genes shared by 3-4 orders or by species representing non-sister orders are most parsimoniously inferred to have been present in the NFNC MCRA. Genes found in the species representing two sister orders do not provide evidence for being present in the NFNC MRCA; they could equally parsimoniously have been present in the NFNC ancestor and lost in the ancestor of the other two orders, or have originated in the ancestor of the sister orders with the gene. Genes found in only a single species are inferred to have arisen at some point in the lineage leading from the species backward to the ancestor of its order.

**Figure 5.**
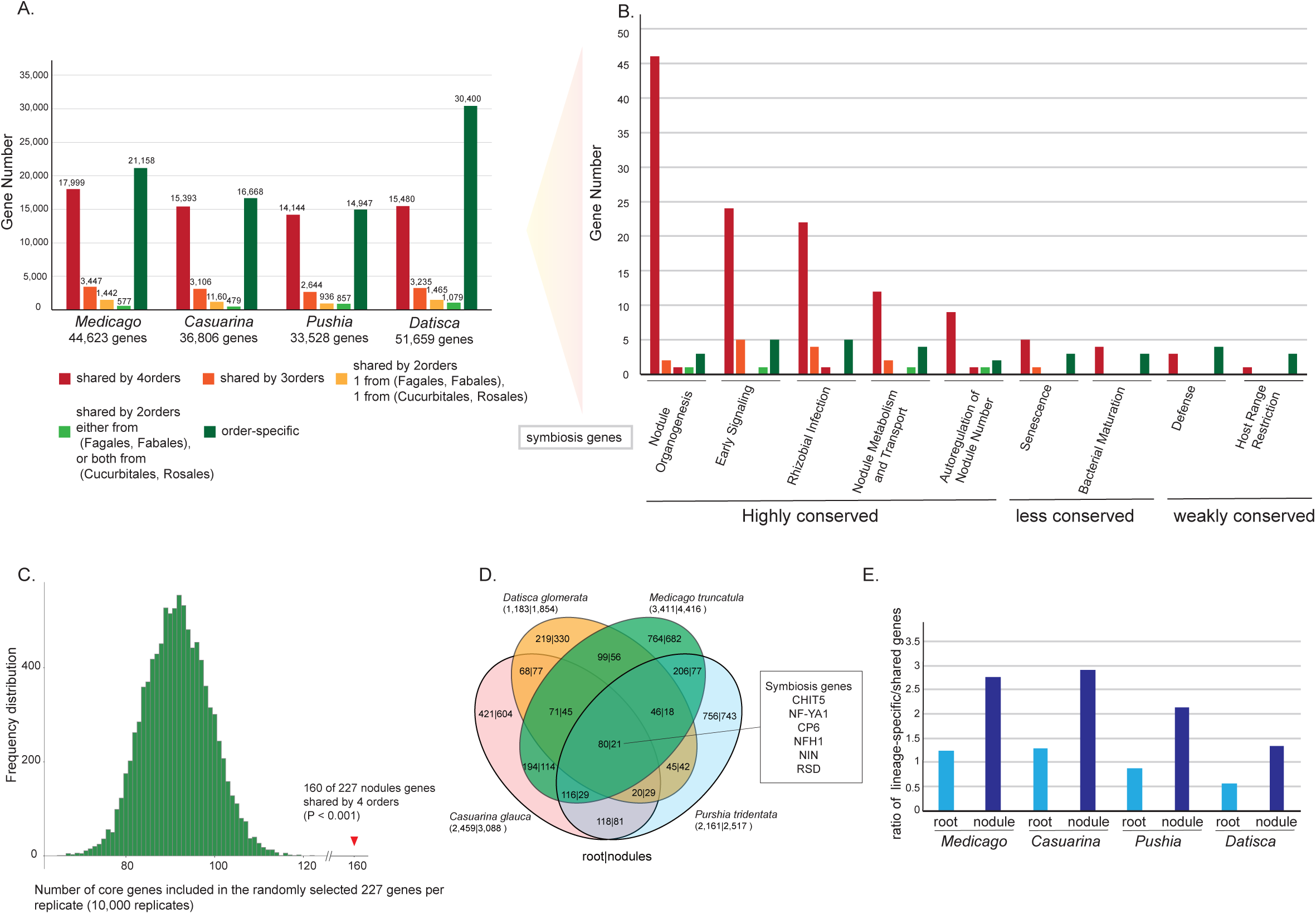
A. identification and catalogue of the shared and lineage-specific genes among the 4 representative nodulating plants in the context of ((Fagales, Fabales), (Cucurbitales, Rosales)). Five categories were analyzed: shared by all 4 species; shared by 3 species; shared by 2 orders, 1 is from the (Fagales, Fabales), 1 is from the (Cucurbitales, Rosales); shared by 2 orders, but either both from (Fagales, Fabales), or both from (Cucurbitales, Rosales); and lineage/order-specific genes. The first three categories are inferred to be present in the most common ancestor of NFNC. The latter two categories arose later than the NFNC MRCA or were lost from either an ancestor of two orders or in the descendants of a single order. B. Evolutionary characterization of genes involved in different aspects of nodulation symbioses as defined in Roy et al. (2020, Fig. 2). Genes are categorized as in Fig. 5A; Highly conserved categories are those for which the ratio of core/lineage-specific genes is >= 4.0; less conserved categories show a lower core/lineage-specific ratio; and weakly conserved categories are those for which lineage-specific genes outnumber core genes C. Frequency distribution of the number of core genes included in the randomly selected 227 genes (10,000 replicates), 160 out of the 227 nodulation-related genes are shared by 4 orders species. D. The distribution of the up-regulated Orthologous Groups (OGs) shared or uniquely present in nodule or root between the 4 selected nodulating plants. Number within Venn plot represent the number of OGs. Number below species name represents the total number of nodule upregulated genes. Numbers on the left side of vertical bar represent root upregulated data, and numbers on the right side represent nodules upregulated data. E. Data from Venn diagram in Fig. 5D, number of DEG+ genes shared among different species. Goal: Calculate the ratio of lineage-specific/shared genes, where “shared” = genes inferred to have been present in the MRCA of the NFNC.

We next determined to which of these categories the 227 *Medicago* nodule symbiosis-related target genes (Roy et al., 2020) belonged. For each of the nodulation processes (e.g., early signaling, nodule organogenesis) into which Roy et al. (2020) divided these genes, we calculated the ratio of core/lineage-specific genes. Most categories were found to be dominated by core genes (**Figure 5B**), despite the fact that lineage-specific genes comprise a high percentage of each of the four genomes. This was particularly true for genes involved in nodule organogenesis, followed by early signaling, rhizobial infection, nodule metabolism and transport, and autoregulation of nodulation (AON). Total gene numbers are much smaller for other categories, but lineage-specific genes dominated the defense and host range restriction categories (**Figure 5B**). Overall, more than 70% (160) of the 227 nodulation-related genes (**Table S2**) are core genes, significantly more than the expectation based on randomly selected sets of 227 genes from the *Medicago* gene set (**Figure 5C**).

We then categorized differentially expressed genes (DEGs) between nodules and roots, and between different lineages (**Table S14-22**). For each species, the ratio of lineage specific to genes likely to have been present in the NFNC MRCA was higher for nodule DEG+ genes than for either nodule DEG- or non DEG genes (**Figure 5E**). Moreover, in all four species, this ratio was approximately double for nodule DEG+ genes than for root DEG+ genes (**Table S28, Figure 5E**). Taken together, these results suggest that recently-evolved genes play a greater role in nodulation than in root biological processes. DEG- and non-DEG lineage-specific genes represent a baseline for genes that have arisen in each order that are not associated with nodulation, so the excess of DEG+ lineage-specific genes suggests that many of these genes could have evolved through involvement in nodulation.

### Conserved non-coding elements (CNEs) associated with nodulation

Recent studies have identified *cis*-regulatory elements that play important roles in the spatio-temporal expression and regulation of symbiosis-related genes (Liu et al., 2019; Soyano et al., 2019). Here, we performed an extensive identification and characterization of CNEs across 88 phylodiverse genomes, by combining two pipelines: 1) the reference-based Lastz-ChainNet/ROAST followed by CNSpipeline (Liang et al., 2018); 2) the reference-free Progressive Cactus-based followed by PhastCons (Hubisz et al., 2011). Our approach differed from that of the recent study of Pereira et al. (2022) by integrating the two pipelines, combining all 88 species for whole-genome alignments followed by CNE identification (**Figure S2**), thereby reducing false positives incorrectly defined as ‘conservation’.

We selected, in addition to *Medicago truncatula*, eight representative genomes comprising one non-nodulator and one nodulator from each of the four orders as well as one outgroup plant (*Populus trichocarpa*) (**Figure S12**) and anchored all the identified CNEs with the *Medicago* genome for comparison. In total, we predicted 931,452 high-quality putative CNEs (>=5bp; corresponding to 4.16% of the *Medicago* genome) **(Figure 6A, Table S29)**, which is a much higher number than that detected by Pereira et al. (2022) (**Figure S13**), who only reported 6,729 CNEs. Furthermore, 84.49% of RNS-specific CNEs and 84.60% of NFN-specific CNEs detected by Pereira et al. (2022) are also present in our study, including the five experimentally validated CNEs in cis-regulatory regions (**Figure S13**), suggesting a high-quality CNE dataset produced as well as many new CNE candidates first discovered in our study (**Figure S2, S13**). As expected, a large proportion (78.4%) of CNEs were located within 20 kb upstream of the 5’ UTR, within 20 kb downstream of the 3’ UTR, or in introns **(Figure 6A, Table S29)**, and many are transposon-related sequences, indicating a rich source of CNEs derived from TEs (**Table S34**). Among the 931,454 CNEs, 331,153 are NFNC-specific, of which 15,101 are RNS-specific. Notably, 3,788 out of the 7,651 nodulation-related CNEs associated with the 232 symbiosis-related genes of Roy et al. (2020; 227 protein coding genes plus 5 miRNA genes) are NFNC-specific (**Table S30-33**). This number is significantly larger than for randomly selected sets of 232 genes (**Figure 6B**), consistent with the hypothesis that at least some CNEs play roles in root nodule symbiosis (Pereira et al., 2022), a trait confined to the NFNC.

**Figure 6.**
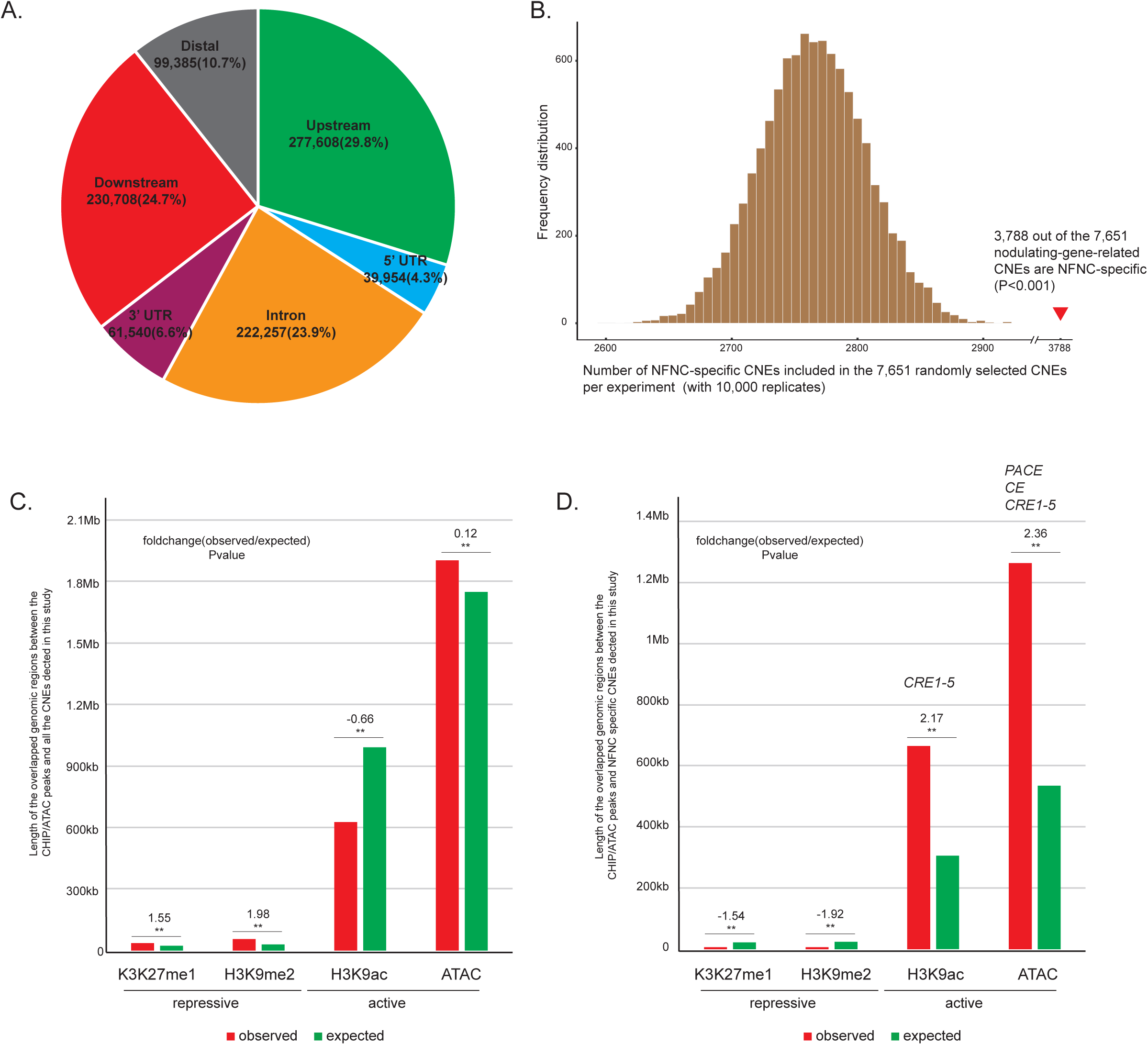
The identification, catalogue, and the evolutionarg and functional analyses of the conserved non-coding elements (CNEs) in the NFNC clade. **A.** Summary statistic on the distribution of CNEs identified by whole-genome alignments across 88 genomes (see Method), with the Medicago truncatula genome as reference. The definition of some genomic elements are given here: distal region, 20kb away from transcriptional start site; upstream, 20kb upstream of the gene; downstream, 20kb upstream of the transcriptional stop site. **B.** Frequency distribution of the number of NFNC-specific CNEs out of the 7,651 randomly selected CNEs for each simulated experiment, with 10,000 replicates in total. The red triangle indicates the number of NFNC-specific CNEs out of the 7,651 nodulation-gene-related CNEs. **C.** Comparison between the genomic regions with the histone markers/ATAC peaks and all the CNEs detected in this study. Significance was calculated by GAT (** P < 0.001). The CHIP/ATAC dataset was downloaded from https://medicago.toulouse.inra.fr/MtrunA17r5.0-ANR. Expected: the frequency distribution of the length of the genomic overlapping regions with the CHIP/ATAC peaks with our simulated experiment in which we randomly selected the same number of CNEs from the whole genome, which were repeated10,000 times. **D.**The same as in C, but we only compared with the NFNC-sepcific CNEs. As control, those CNEs with functional validation, functional NFNC specific CNEs, PACE, CE and CRE1-5 were overlapped with ATAC peak, and CRE1-5 also overlap with H3K9ac peak.

To further evaluate the quality and function of the CNEs identified in this study, we compared each category of CNEs with the PLACE and plantPAN3.0 database (**Figure S2B**). We calculated the enrichment of motifs based on Z-score for each CNE and retrieved their hits from the databases, and found that 52.94% of CNEs have at least one hit in the database, suggesting the potential functional roles of these CNEs. Notably, one nodulation-related motif ‘AAAGAT’, originally identified in the soybean leghemoglobin *lbc3* promoter (Fehlberg et al., 2005; Stougaard et al., 1990), ranks in the top 10 sorting by the Z-score based on the CNEs that were conserved in all nodulating plants (**Figure S2B**). At least 4,468 CNEs contain the ‘AAAGAT’ motif located around symbiosis genes, suggesting that an amplification of this motif in nodulating plants might have contributed to root nodule symbiosis emergence (**Figure S2B**).

The distribution of CNEs in the *Medicago* genome was compared (separately for all 931,454 CNEs and for only the 331,153 NFNC-specific CNEs) with various features associated with regulation of transcription (two heterochoromatic repressive histone marks, H3K9me2 and H3K27me1; one active mark, H3K9ac; and open chromatin as assayed by ATAC-seq). (**Figure 6C, D**, and **Figure S9**). CNEs were enriched at the H3K9ac marks around transcriptional start and end sites in both nodules and roots, which could promote the expression of their associated gene **(Figure S10).** Interestingly, we found that a significant majority of the NFNC-specific CNEs overlap and are enriched with the active marker H3K9ac and the active ATAC-seq peaks (**Figure 6D**), suggesting that some of the NFNC-specific CNEs play roles in activating the initiation of genes involved in root nodule symbiosis.

Two legume specific *cis*-regulatory elements were detected in our study. One is a remote cis-regulatory region located 20 kb upstream of the translational start site of *NIN* (**Figure S10**). This is consistent with the fact that the putative cytokinin-responsive elements within this region are required for the triggering of *NIN* expression in the pericycle by cytokinin and are indispensable for nodule organogenesis (Liu et al., 2019). Interestingly, the other legume-specific *cis*-regulatory element is a NIN-binding site located in the intron of *LBD16a* (**Figure S11**). *LBD16*a is involved in both lateral root formation and nodule organogenesis, of which only the latter is dependent on NIN (Soyano et al., 2019).

## Discussion

For many years it has been accepted that nodulation is a complex phenomenon, many of whose components were recruited from pre-existing processes that are shared widely among nodulating and non-nodulating plants, both within and outside of the single angiosperm clade, NFNC, in which nodulating species occur. What is unique about nodulation is therefore the assembly of these disparate components into a functional association in which bacteria that would otherwise be pathogenic are attracted, their entry past highly effective organ- and cell-level defenses is facilitated, existing developmental programs are co-opted to build a home for them, nutrition is provided, and the nitrogen-containing fruits of their labor are harvested and transported into the plant. How did this assembly process occur? Were there intermediate states, analogous for example to the origin of key features of avian wings prior to their use in flying (Uno and Hirasawa, 2023). If so, might some extant non-nodulating species be proto-nodulators? Finding examples of non-nodulating species that have some but not all components of nodulation would be an exciting development in efforts to engineer symbiotic nitrogen fixation outside of the NFNC, and argues for investment of research on the non-nodulating relatives of nodulators across the NFNC. Such plants would represent steps toward nodulation under the multiple origins Scenario II, or plants that had lost the full ability to produce nodules, but for which cessation of nodulation had not led to the loss of all nodulation-related features, in Scenario I. The latter case would be of particular interest if loss was avoided because some selective benefit was still provided by the retained features.

Developing tests that distinguish between homology and convergence of nodulation is very difficult due to the complexity of deep homology, itself an evolving concept in evolutionary theory, with recent emphasis on combinatorial “character integration mechanisms” as the generators of evolutionary novelty (DiFrisco et al., 2022). As such, it is possible that there are no genes or GRNs truly unique to nodulation. In any case, “unique” is a relative term when so much of gene evolution involves duplication and neo- or subfunctionalization, and given the complex and often overlapping roles that individual genes perform such that a high percentage of the genome is expressed even in single cell types (e.g., (Coate et al., 2020). Here we have increased genomic and transcriptomic sampling of phylogenetically diverse nodulating and non-nodulating taxa in order to identify shared and divergent elements of nodulation. Even after our efforts to identify other key genes, the transcription factor, *NIN*, remains the best candidate for a gene specifically associated with nodulation. As our results show, much remains to be learned about its structure and function across the NFNC, and achieving a greater understanding of this key gene will be important regardless of whether nodulation evolved once or multiple times.

A major debate in evolutionary biology concerns the relative contributions of novel regulatory sequences versus protein coding genes to evolutionary innovation (Carroll, 2005; 2008; Stern, 2000). In the case of nodulation, specific *cis*-regulatory element changes in *LjLBD16* and *MtSCR* have been shown to be required for legume-rhizobial symbiosis (Dong et al., 2021). Recruitment of genes for nodulation likely involved the evolution of new regulatory sequences associated with genes that also retained their original functions, making characterization of such sequences critical to understanding not only the process of recruitment—including potentially distinguishing between the two evolutionary scenarios (Doyle, 2016) —but also how gene regulatory processes are modified when nodulation is added to existing developmental programs. Conserved noncoding elements (CNEs) include regulatory elements (e.g., (Schmitz et al., 2022), and a recent study has identified many such elements associated with nodulation, and validated one such element experimentally (Pereira et al., 2022).

Here, we developed a novel pipeline to identify CNEs, combining the merits of the Lastz-based and the Cactus-based pipelines, to identify a much larger set of NFNC-specific and RNS-specific CNEs. We identified thousands of non-coding sequences that are highly conserved among nodulating species, many of which are associated with symbiosis-related genes. This provides a vast pool of candidates in the search for nodulation-associated regulatory elements (Pereira et al., 2022), including those currently associated with nodulation and those that could provide fossil evidence of former nodulation ability in non-nodulating species (Doyle, 2016). However, CNE identification and characterization is still a challenging task, requiring additional high-quality chromosome-level genome assemblies and annotation, as well as functional validation for the identified conserved non-coding elements in the context of regulatory genomics (e.g., Pereira et al., 2022).

The phylogenetic diversity of nodulating species provides an opportunity to explore the many different solutions these lineages have evolved for attracting and housing bacteria. The question of whether the various modules were assembled once or many times, as fascinating as it is, pales in significance compared to determining in detail how nodulation can produce the same result in taxa some of which diverged over 100 MYA, involving diverse bacteria housed in structures that are highly divergent despite developmental commonalities. If Scenario I is correct, then differences in how nodulation occurs in such lineages provides information on the robustness of an ancestral symbiosis. If Scenario II is correct, then convergent similarities represent the requirements for establishing a nodulation symbiosis de novo. In either case there is a clear need for additional phylogenomic and phylotranscriptomic sampling and deep comparative biology analysis for both protein-coding genes, gene expression, and conserved non-coding elements across the entire NFNC, as proposed in The Legume Nodulation and NFNC Phylogenomics v2.0 Project (https://www.legumedata.org/beanbag/68/issue-68-legume-genome-sequencing-consortium).

## Materials and Methods

### Plants and bacteria

Seeds of *Alnus glutinosa* (harvested from a tree growing on the Rhône River banks inin Lyon, France) and *Betula pendula* (Vilmorin, La Ménitré, France) were sowed, left to germinate and grown for six weeks in a sterile soil/vermiculite substrate (1:1, vol/vol) in a greenhouse with a 16 h light/8 h dark regime with temperatures of 21 °C (light) and 16 °C (dark). Seedlings were transferred to 500 ml Fåhraeus medium (Fåhraeus, 1957) with or without 5 mM KNO_3_ in opaque plastic pots (8 seedlings per pot) and grown for four weeks before inoculation (or not). Inoculation was performed using syringed 18-day-old *Frankia alni* ACN14a culture in BAP-PCM medium containing 5 mM NH_4_Cl pH 6.2 (Schwencke, 1991). Four treatments were thus done: 1. seedlings with 5 mM KNO_3_ and *Frankia*, 2. seedlings with 5 mM KNO_3_ without *Frankia*, 3. seedlings without KNO_3_ and with *Frankia*, 4. seedlings without KNO_3_ and without *Frankia*. After inoculation, plants were grown for 22 days. Then, roots – nodulated or not – were harvested and frozen in liquid nitrogen.

Cuttings of *Begonia fuchsioides* were obtained from the Nymphenburg Botanical Garden in Munich (Germany) in 2015 and grown in a growth cabinet at low light (2 fluorescent lights removed from the cabinet) at 16 h light (18.5 °C) / 8 h dark (12 °C). They were grown in a 1:1 (v/v) mixture of sand (grain size 1-1.2 mm) / Stender Vermehrungssubstrat A 210 (Stender AG, Schermbeck, Germany). Plants were regularly watered with deionized water and once a week with ¼ Hoagland’s (Hoagland and Arnon, 1950) – either without nitrogen (minus N samples) or either containing 10 mM KNO_3_ (plus N samples). Material from cuttings was harvested after *ca.* three months of growth, and shock-frozen in liquid nitrogen. Roots and leaves were frozen separately. The entire root systems were harvested except for the top part of ca. 1 cm which showed secondary growth and lignification.

*Casuarina glauca* Sieb. Ex Spreng seeds were purchased from the Australian Tree Seed Centre (CSIRO, Australia) and grown as described by (Auguy et al., 2011). The compatible bacterial strain *Frankia casuarinae CcI3* (Zhang et al., 1984) was used to inoculate *C. glauca* plantlets as previously described (Franche et al., 1997). Seedlings were transferred to a soil/vermiculite substrate (4:1, vol/vol) in a greenhouse under natural light at temperatures between 25 and 30°C. After 1 month, seedlings were transferred to pots containing 500 ml of a modified Broughton and Dillworth (BD) (Broughton and Dilworth, 1971) medium supplemented with nitrogen (5 mM KNO_3_) and cultivated in a growth chamber under the following conditions: 25°C, an average 45% humidity and a 16-h photoperiod, Photosynthetically Active Radiation (PAR) of 150 µmol m^-2^ s^-1^. After 3 weeks, plants were starved of nitrogen for 1 week prior to inoculation with the symbiotic bacteria. Plants were inoculated with 10 ml of a concentrated *Frankia casuarinae* CcI3 suspension at a density corresponding to ∼25 μg ml^−1^ of protein (Franche *et al*., 1997). After 2 h of contact, plants were placed in pots containing 490 ml of BD without nitrogen and 10 ml of the CcI3 suspension. Nodule initiation was monitored twice per week. Six conditions were sampled: leaves, non-inoculated roots with or without KNO_3_, inoculated roots (2, 4, 8 days after inoculation) and 3 weeks old nodules. All samples were immediately frozen in liquid nitrogen.

*Datisca glomerata* (C. Presl) Baill. seeds originating from plants growing at Gates Canyon in Vacaville, California, USA, were brought to Europe in 1995 by Katharina Pawlowski and propagated in greenhouses ever since. For roots, plants were grown in axenic culture. Seeds were surface-sterilized by incubation in 25% H_2_SO_4_ for 30 s, followed by two washes with sterile deionized water. Then they were incubated for 5 min in 2.5 % NaOCl and washed six times with sterile deionized water. Seeds were transferred to vertical Petri dishes with ¼ Hoagland’s with 10 mM KNO_3_ and 1% agar, or to vertical Petri dishes with ¼ Hoagland’s without nitrogen and 1% agar. Roots were harvested after 7 weeks of cultivation. For nodulation, seeds were transfered to the greenhouse to pots with germination soil (Såjord, Weibull Trädgard AB, Hammenhög, Sweden) covered by sand (1.2-2 mm Quartz; Radasand AB, Lidköping, Sweden). Greenhouse conditions were 13 h light (23 °C) / 11 h dark (19 °C) and 200 µmol m^-2^ s^-1^ PAR. When seeds had germinated, seedlings were transferred to little pots with germination soil. For infection with *Candidatus* Frankia datiscae Dg1 (Persson et al., 2011), plantlets of 10 cm height were transferred to larger pots (diameter 15 cm) containing a 1 :1 (v/v) mixture of sand (0–2 mm Quartz; Rådasand AB, Lidköping, Sweden) and germination soil (bottom third), sand (middle third) and an 1:1 (v/v) mixture of sand and germination soil (top third). Inoculum was applied to the root system during transfer in the form of *D. glomerata* nodules, freshly harvested from an older inoculated plant and crushed in deionized water with mortar and pestle. Inoculated plants were watered with ¼ Hoagland’s without N once per week, otherwise with deionized water. Nodules were harvested 6-12 weeks after infection. For leaf production, *D. glomerata* seeds were germinated after two weeks of vernalization, and then transferred to pots and grown in the greenhouse at LMU Munich (18 °C/12 °C, 16 h/8 h day/night cycles, 150 μmol m^-2^ s^-1^ PAR). Growth substrate was A210 (perlite, with 0.5 kg/m3 of 14-10-18 NPK, with Sphagnum peat H3-H5, pH 6.2) from Stender AG (Schermbeck, Germany). Plants were regularly watered with deionized water and once a week with ¼ Hoagland’s with 10 mM KNO_3_. Leaves were harvested after 6-12 weeks of growth.

*Purshia tridentata* seedlings were purchased in 2012 from Cornflower Farms Native Nursery, in Elk Grove, CA (USA), where they had been grown in soil mix with slow-release nitrogen fertilizer. The nursery soil mix was replaced with pasteurized soil mix, sand:fir bark:peat moss:perlite. Seedlings were maintained on deionized water (DI) before and after inoculation. At one week post-transplant, the seedlings were inoculated with an aqueous suspension of rhizosphere soil that was excavated from mature shrubs of *Ceanothus velutinus* at Sagehen Creek Field Station, Truckee, CA (USA), and stored at 4°C. Nodulation was observed at 95 days post-inoculation. Thereafter, the nodulated *P. tridentata* plants were maintained as stock plants in greenhouse conditions at the University of California, Davis, CA (USA), and were irrigated with DI, except for two supplements of Hoagland’s solution, 10 ml of ½-strength Hoagland’s/5L container (23/09/2014; 26/02/2015). The ½-strength Hoagland’s in the greenhouse contains 150 mg N/L. Thus the 10-ml supplement was equivalent to 1.5 mg N/5L container. In July, 2015, to test whether recent application of nitrogen affected nodule-lobe or root-tip gene expression, one-half the plants from which nodules and roots were collected were given two supplements of Hoagland’s solution, 20 ml of 1/2-strength Hoagland’s per 5 L container; while the other one-half of the plants used for sampling did not receive any supplement. Twenty ml of the greenhouse ½-strength Hoagland’s solution (150 mg/L N) is equivalent to 3 mg N/5L container Samples collected from the *P. tridentata* root systems consisted of mature nodule lobe tips (i.e. the portion of individual perennial nodule lobes from the current growing season, that had reached their full extent for the season) and root tips (< 4cm length). Nodule lobe tips and root tips were rinsed in sterile deionized water, immediately frozen in liquid nitrogen, and stored at −80° C.

### RNA isolations

For *A. glutinosa* and *B. pendula,* RNA was extracted as described by (Alloisio et al., 2010) using the RNAeasy Plant Mini Kit (Qiagen) and on-column DNA digestion with the RNase-free DNase set (Qiagen). In order to remove any remaining DNA contamination, a second DNase treatment was performed with RQ1 RNase-free DNase (Promega, Charbonnières-les-Bains, France), followed by RNA clean-up using the RNeasy Plant Mini Kit. Purity, concentration and quality of RNA samples were checked using a NanoDrop 1000 spectrophotometer (Thermo Fisher Scientific, Courtaboeuf, France) or an Ultrospec 3300 Pro photospectrometer (Amersham Biosciences, Buckinghamshire, UK) and agarose gel electrophoresis.

For *C. glauca*, two conditions were sampled within three biological replicates: 21 days old nodule and roots were sampled from inoculated and non-inoculated plants (control), respectively. Total RNA was purified by ultracentrifugation (Hocher et al., 2006). Residual DNA was removed from RNA samples using the Turbo DNA free kit (Ambion, USA), quantified using a NanoDrop ND-1000 spectrophotometer (Thermo Fisher Scientific, Courtaboeuf, France). We used a pool of 28 plants for each time point. The integrity of the RNA samples was assessed using a Bioanalyzer 2100 according to the manufacturer’s instructions (Agilent, Santa Clara, CA, USA). mRNA libraries were constructed for each condition and sequencing was performed at the MGX platform (Montpellier Genomix, *Institut de Genomique Fonctionnelle*, Montpellier France). The RNA libraries were constructed using the TruSeq stranded mRNA library construction kit (Illumina Inc., USA). The quantitative and qualitative analyses of the library were carried on Agilent DNA 1000 chip and qPCR (Applied Biosystems 7500, SYBR Green). RNA was sequenced using the Illumina SBS (sequence by synthesis) technique on a Hiseq2000 in single read 100 nt mode. Image analysis, base calling and quality filtering were performed using Illumina software.

For *D. glomerata* and *B. fuchsioides,* RNA isolations were performed using the Sigma Spectrum^TM^ Plant Total RNA extraction kit (Sigma-Aldrich, St. Louis, MO, USA); Polyclar AT (Serva, Heidelberg, Germany) was added to the extraction buffer (2% w/v). On-column DNA digestion with the RNase-free DNase set (Qiagen), followed by treatment with Ambion TURBO DNase (ThermoFisher) after isolation. RNA quality was determined using a NanoDrop 1000 spectrophotometer (Thermo Fisher Scientific).

RNA from *P. tridentata* samples was extracted using either Qiagen RNeasy Plant Mini-Kit, or for a subset of samples, the Spectrum Plant Total RNA Kit, according to instructions in the kits, followed by treatment with Ambion Turbo RNAse-free DNAse kit (Thermo Fisher Scientific) after isolation. The Spectrum Total RNA kit was used to enable BGI to test whether *Frankia* transcripts could be detected in nodule tissue, in addition to plant gene expression.

### Genome sequencing, assembly, and annotation

Plant growth, DNA extractions, and RNA extractions were carried out following the protocol described in Griesmann et al., 2018 (Griesmann *et al*., 2018). The genomes of *Purshia tridentata, Dryas octopetala* and *Fagus sylvatica* were sequenced using Illumina sequencing technology (Hiseq2000 and Hiseq4000). A hierarchical DNA library strategy was employed, which included multiple paired-end libraries with insert sizes ranging from 170-800bp and mate-pair libraries using large DNA fragments with insert sizes of 2kb-20kb. The reads were filtered using SOAPfilter (Luo et al., 2015) following a strict quality control protocol. The genome size was estimated using 17-mer analysis (Luo *et al*., 2015).

Whole genome assembly was performed using SOAPdenovo2 (Luo et al., 2015), ALLPATH-LG (Luo et al., 2015), and Platanus (Kajitani et al., 2014). Repetitive elements were identified and analyzed using RepeatMasker (Bergman and Quesneville, 2007) and RepeatModeler (Flynn et al., 2020).

The MAKER-P pipeline (Campbell et al., 2014) was used for gene annotation by integrating multiple annotation resources. The genome annotation revealed 23155, 28191, and 35140 genes in the genomes of *Purshia*, *Dryas*, and *Fagus*, respectively. Gene functional prediction and assignment were performed using Interproscan (Jones et al., 2014) and homologous searches against the Swiss-Prot database (https://www.ebi.ac.uk/uniprot) and the KEGG database (https://www.kegg.jp/blastkoala/).

### Evolutionary analysis

#### Analysis of Reciprocal Best Hits (RBHs) and orthologous/paralogous group

The genome sequences and annotations of the selected plants in this study were downloaded from NCBI (https://www.ncbi.nlm.nih.gov/) and other plant genome websites (Table S1). To elucidate the orthology of all genes, we used two strategies - identifying Reciprocal Best Hits (RBHs) and orthologous/paralogous clustering. First, we identified RBHs by performing all-vs-all BLAST searches of the plant proteomes against the *Medicago truncatula* proteome, and then performing reverse-BLAST searches of the Medicago proteome against the other plant proteomes. We analyzed the presence and absence of the orthologs of RBHs for each species. We compared the number of species carrying orthologs of target reference proteins between nodulators, non-nodulators, and outgroups using Fisher’s exact test to infer the association between the presence of target genes and nodulation. We calculated p using the formula:

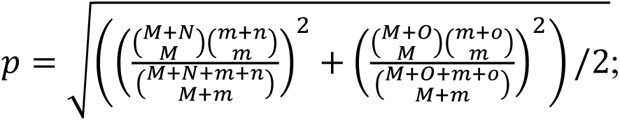

Here, M, N, and O are the numbers of nodulators, non-nodulators, and outgroup species, respectively, carrying orthologs of target proteins; m, n, and o are the number of nodulators, non-nodulators, and outgroup species, respectively, not carrying orthologs of target proteins. Second, we performed all-vs-all BLAST searches of all plant proteomes and used Orthofinder to identify the orthologous/paralogous groups (Emms and Kelly, 2019). We calculated the gene copy number of each group in each species. We then compared the copy numbers of protein family members from nodulating (legume or non-legume), non-nodulating species, and outgroups using a *T*-test. If the difference of copy number revealed by Orthofinder between two groups of interest (i.e., nodulating species vs. non-nodulating species) for a gene family is larger than one and the t-test value smaller than 0.01, that family was considered as an expanded gene family in nodulating species. To confirm the orthology of families revealed by these two strategies, we constructed the phylogenetic tree of each target gene. We aligned the proteins encoded by target genes from each species using MAFFT (Katoh et al., 2002), and then constructed maximum-likelihood phylogenetic trees with the best-fit model selected by IQ-tree and bootstrap 1000 (Nguyen et al., 2015).

#### Identification of convergent loci/sites

We used two combinatorial approaches to detect convergent signals within sequences in nodulating species: a ‘tree topology inference’ method and an ‘alignment’ method.

For the ‘tree topology inference’ method, we used a pipeline based on maximum likelihood (ML) phylogenetic reconstruction according to the Parker et al. method (Parker et al., 2013; Thomas and Hahn, 2015) with minor modifications. We aligned each orthologous protein along with CDS and measured its fit (site-wise log-likelihood support; SSLS) to the known species tree (H0) and an alternative topology in which nodulating species (H1) formed a monophyletic clade. The pipelines were briefly described as follows:1) the one-to-one orthologs present in all nodulators or non-nodulators were aligned using MAFFT; 2) the log-likelihood of phylogenies (H0, H1) was calculated for every site in the alignment using the RAXML (v8.2.12) program (Stamatakis, 2014) with the parameters set as “-f g -m GTRGAMMA ”; the resulting log-likelihoods of H0 and H1 for each site were subtracted to generate the ΔSSLS as: ΔSSLSi = lnLi,H0 – lnLi,H1, where lnLi,H0 and lnLi,H1 denote the log-likelihood of the ith site under H0 and H1, respectively; the sequence convergence of each gene was quantified by taking the mean of ΔSSLS at each sites; 3) ten thousand random trees were generated to simulate the null distribution of ΔSSLS. ΔSSLS of a significant convergent gene (support H1) should be smaller than left tail 0.01 probability of the null distribution. Since a few key nucleotides/residues may alter the function of protein, we next used the alignment method to detect convergent sites. For the ‘alignment’ method, convergent sites were detected by a perl script that searched for similar amino acids present in all nodulating species but not in non-nodulating species based on the alignment of one-to-one orthologs, where the similar amino acids were defined as the BLOSUM score > 0 for pairwise comparison (codes of the pipeline are available on https://github.com/yu-z/NFN-phylogenomics).

### Identification of conserved non-coding elements

We identified Conserved Non-coding Elements (CNEs) by comparing the *M. truncatula* reference genome with 87 query genomes (**Figure S2, Figure S13**) (Armstrong et al., 2020; Haudry et al., 2013; Hubisz *et al*., 2011; Liang *et al*., 2018; Sackton et al., 2019). First, we annotated simple repeats using Tantan (Frith, 2011) to find orthologous sequences more accurately. Multiple sequence alignments of whole genomes were generated separately using two methods named lastz-chainNet-roast (Liang *et al*., 2018) and Progressive Cactus (Armstrong *et al*., 2020). For the lastz-chainNet-roast method, each query genome was aligned to the *Medicago* genome using LASTz (v1.04.00) (Harris, 2007) with parameters “--ambiguous=iupac --chain --notransition H=2000 Y=3000 L=3000 K=2200 --format=axt --gfextend”; the alignments for each query were linked into longer chains using axtChain, chainPreNet and chainNet with default parameters (Kent et al., 2003). For each query, when two alignments overlapped in the *Medicago* genome, the overlapping part of the shorter one was removed by single_cov2 (http://www.bx.psu.edu/~cathy/toast-roast.tmp/README.toast-roast.html). Next, we linked all the pairwise alignments of each query genome and the *Medicago* reference genome according to the topology of the species-tree using ROAST (v3; http://www.bx.psu.edu/~cathy/toast-roast.tmp/README.toast-roast.html). For the Progressive Cactus method, the one-to-one orthologs phylogenetic tree was used as guide tree and the high-quality assembly (scaffold N50 >=1 Mb and contig N50 >= 20 Kb) were marked with *. The reference-free alignment in HAL format was export to maf alignment using different species (*Medicago truncatula*, *Ammopiptanthus nanus*, *Discaria trinervis*, *Ziziphus jujuba*, *Datisca glomerata*, *Lagenaria siceraria*, *Casuarina glauca*, *Fagus sylvatica*, *Populus trichocarpa*) as references. Maf alignments using *Medicago truncatula* as reference from two methods were combined together. Only alignments with length ≥ 5bp and rows ≥ 10 were retained. PhastCons (Hubisz *et al*., 2011) and CNSpipeline were used separately to identify conserved elements (CEs). For PhastCons, we use default parameters to identify CEs. For CNSpipeline, we calculated the conserved score for each alignment: score = number of aligned sequences / total number of interested species. Alignment with score >= 0.9 were selected as conserved CEs in each species. Then, we merged CEs generated by two CEs identification methods (merged overlap CEs and remained specific CEs). CEs which only overlapped with non-coding region were referred as conserved non-coding elements (CNEs). The presence/absence analysis of CNEs was performed the same way as the presence/absence analysis of genes above. The overlappeing CNEs between nodulating and non-nodulating species were calculated by a series of Python Scripts. The significance of enrichment of motifs of CNEs were calculated by a 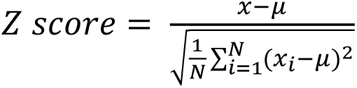, where x_i_ is the number of CNEs that matched a certain not less than 6nt motif in the PLACE and plantPAN3.0 (http://plantpan.itps.ncku.edu.tw/) database (*x*_*i*_ ≥ 2), N is the total number of matched motifs, *μ* is the average of number of CNEs matched motifs. To investigate the enrichment of two repressive histone marks (H3K27me1 and H3K9me2), one activating mark (H3K9ac) and ATAC data in CNEs, GAT (Heger et al., 2013) were used to compare the genome locus of CHIP mark and CNEs.

### Hairy Root Transformation

The CE region (Liu *et al*., 2019), 5-kb promoter, and full-length CDS of wild-type *LjNIN* and its variants were cloned into pUB-GFP vector to obtain pUB-GFP LjNINproCE-LjNIN-wt pUB-GFP LjNINproCE-LjNIN-TA, and pUB-GFP LjNINproCE-LjNIN-TD, respectively. *A. rhizogenes* strain LBA1334 cells carrying pUB-GFP LjNINproCE-LjNIN-wt pUB-GFP LjNINproCE-LjNIN-TA, and pUB-GFP LjNINproCE-LjNIN-TD, and empty vector were used to induce hairy root formation in *Ljnin-2* mutant plants (*Lotus japonicus*) using a procedure as described previously (Yuan et al., 2012). Phenotypes of transgenic hairy roots were screened and photographed 21 days after inoculation with *M. loti* MAFF303099. Transgenic hairy roots expressing the empty vector (pUG-GFP) were used as a negative control. The mean values of nodule number and Student’s t-tests were performed using R.

### RNA-seq analysis

We designed systematic meta-analysis of transcriptome on a variety of species from NFNC. We obtained 151 RNA-seq libraries across 20 phylodiverse species within the NFNC, among which 7 species were highlighted for comparison across tissues (roots/nodules) and treatments (with or without nitrate and/or compatible *Frankia*), including 4 actinorhizal nodulating plant species (*Alnus glutinosa, Datisca glomerata, Casuarina glauca, Pushia tridentata)* and 2 non-nodulating species (*Betula pendula, Begonia fuchsioides)* (**Figure 4A, Table S15-27**). We added 9 transcriptomes of *Medicago truncatula* from NCBI for comparative analysis.

The raw RNA-seq reads of low-quality were filtered by soapfilter (Luo *et al*., 2015). For species of which genome have been sequenced, we mapped filtered reads to the reference genome using HISAT2 (Kim et al., 2019). Reference guided transcript assembling for each libraries was executed using StringTie (Pertea et al., 2015). Assembled transcripts from each library were then merged, and the transcripts on the genome was re-annotated using gffcompare. For species for which the genome has not been sequenced, the transcripts were *de novo* assembled using Trinity (Grabherr et al., 2011). We calculated the expression level (TPM) of each gene using Salmon (Patro et al., 2017). The pairwise correlations between transcriptomes were calculated by gene expression levels (as estimated by TPM) of 3987 one-to-one orthologs across the 21 species (including the public dataset, Medicago) (**Figure S7**). The genes differentially expressed in different treatments (+N-B, +N+B, -N+B, -N-B) were analyzed by DESeq2 (adjusted *p* value < 0.05 and log2FoldChange > 2) (Love et al., 2014). The ortholog/paralog groups of DEGs upregulated in the same tissue/treatment of different species identified by Orthofinder. The symbiosis genes were mapped to each group, and the variation between species were analyzed by a series of perl Scripts. The differentially expressed genes and the encoded proteins were functionally annotated by interproscan, and Swiss port, BLAST and KEGG (https://www.kegg.jp/blastkoala/).

## Data availability

The raw RNA sequencing data, DNA sequencing data, as well as the new genome assemblies and annotations have been deposited into CNGB Sequence Archive (CNSA) of China National GeneBank DataBase (CNGBdb) with accession number CNP0004055. Multiple whole genome alignment files (Cactus) have been uploaded to Zenodo (https://zenodo.org/record/5798193#.ZCGPt-zP30p).

## Author contributions

S.C. designed and oversaw the study. S.C., J.D. and Y.Z. wrote the manuscript. W.X., Y.F., Y.Z., and X.L., analyzed data. Alison Berry provided samples for *Purshia tridentata*; Pierre-Marc Delaux provided samples of *Mimosa pudica*; Valerie Hocher provided samples of *Casuarina glauca*; Martin Parniske provided samples of *Dryas drummondii* and *Dryas octopetala*. Katharina Pawlowski provided samples of *Begonia fuchsioides* and *Datisca glomerata*. Petar Pujic, Nicole Alloisio, Pascale Fournier, Hasna Boubakri and Philippe Normand provided DNAs of *Alnus glutinosa*, *Betula nana*, and *Fagus sylvatica* and contributed their RNA-seq experiments.

## Acknowledgements

We thank Douglas Soltis and Pamela Soltis for providing suggestions. S.C. is supported by the National Natural Science Foundation of China (32022006), the Program for Guangdong “ZhuJiang” Innovation Teams (2019ZT08N628), the Agricultural Science and Technology Innovation Program (ASTIP) (CAAS-XTCX2016001) and the special funds for science technology innovation and industrial development of Shenzhen Dapeng New District (PT202101-01). Y.Z. is supported by the National Natural Science Foundation of China (32070250), the Natural Science Foundation of Guangdong Province (2020A1515011030) and the open research project of “Cross-Cooperative Team” of the Germplasm Bank of Wild Species, Kunming Institute of Botany, Chinese Academy of Sciences. For Casuarina *glauca* RNA seq data, financial support was provided by IRD and the Agence Nationale de la Recherche (Project SESAM 2010 BLAN 1708 01).

